# Computational optimization of the SARS-CoV-2 receptor-binding-motif affinity for human ACE2

**DOI:** 10.1101/2020.07.20.212068

**Authors:** S. Polydorides, G. Archontis

## Abstract

The coronavirus SARS-CoV-2, that is responsible for the COVID-19 pandemic, and the closely related SARS-CoV coronavirus enter cells by binding at the human angiotensin converting enzyme 2 (hACE2). The stronger hACE2 affinity of SARS-CoV-2 has been connected with its higher infectivity. In this work, we study hACE2 complexes with the receptor binding domains (RBDs) of the human SARS-CoV-2 and human SARS-CoV viruses, using all-atom molecular dynamics (MD) simulations and Computational Protein Design (CPD) with a physics-based energy function. The MD simulations identify charge-modifying substitutions between the CoV-2 and CoV RBDs, which either increase or decrease the hACE2 affinity of the SARS-CoV-2 RBD. The combined effect of these mutations is small, and the relative affinity is mainly determined by substitutions at residues in contact with hACE2. Many of these findings are in line and interpret recent experiments. Our CPD calculations redesign positions 455, 493, 494 and 501 of the SARS-CoV-2 RBM, which contact hACE2 in the complex and are important for ACE2 recognition. Sampling is enhanced by an adaptive importance sampling Monte Carlo method. Sequences with increased affinity replace CoV-2 glutamine by a negative residue at position 493, and serine by nonpolar, aromatic or a threonine at position 494. Substitutions at positions positions 455 and 501 have a smaller effect on affinity. Substitutions suggested by our design are seen in viral sequences encountered in other species, including bat and pangolin. Our results might be used to identify potential virus strains with higher human infectivity and assist in the design of peptide-based or peptidomimetic compounds with the potential to inhibit SARS-CoV-2 binding at hACE2.

**SIGNIFICANCE:** The coronavirus SARS-CoV-2 is responsible for the current COVID-19 pandemic. SARS-CoV-2 and the earlier, closely related SARS-CoV virus bind at the human angiotensin converting enzyme 2 (hACE2) receptor at the cell surface. The higher human infectivity of SARS-CoV-2 may be linked to its stronger affinity for hACE2. Here, we study by computational methods complexes of hACE2 with the receptor binding domains (RBDs) of viruses SARS-CoV-2 and SARS-CoV. We identify residues affecting the affinities of the two domains for hACE2. We also propose mutations at key SARS-CoV-2 positions, which might enhance hACE2 affinity. Such mutations may appear in viral strains with increased human infectivity and might assist the design of peptide-based compounds that inhibit infection of human cells by SARS-CoV-2.

## INTRODUCTION

Coronaviruses are enveloped, single-stranded RNA viruses responsible for respiratory, gastrointestinal and central nervous system diseases in various avian and mammalian species (1). The novel SARS-CoV-2 coronavirus caused acute respiratory syndrome and pneumonia in humans at the end of 2019 in Wuhan, China, and has since caused a pandemic responsible for over 13 million confirmed cases and 500000 deaths (early July 2020). In addition to SARS-CoV-2, other coronaviruses able to infect humans are the severe acute respiratory syndrome virus (SARS-CoV) and the Middle East respiratory syndrome virus (MERS-CoV), which appeared, respectively, in 2002 and 2012, and the mild-symptom viruses HKU1, NL63, OC43 and 229E (2, 3). SARS-CoV-2 is closely related to SARS-CoV and the bat CoV ZC45, RmYNo2, RaTG13, sharing with them a nucleotide sequence identity of 79.5%, 89.1%, 93.3%, and 96.2%, respectively (4–6).

Receptor binding and membrane fusion are key initial steps in the coronavirus infection cycle. Entry into host cells is mediated via the viral spike (S) protein, which forms large protuberances at the virus surface and gives the crown (corona) appearance to the coronaviruses. The S protein (reviewed in (7)) consists of an ectodomain, a transmembrane domain and an intracellular tail. The ectodomain contains a receptor-binding subunit (S1) and a membrane-fusion subunit (S2). During virus entry, the S1 subunit binds to a receptor on the host cell surface and the S2 subunit fuses the host and viral membranes, allowing viral genomes to enter host cells (7).

The host receptor of the human SARS-CoV-2 and SARS-CoV S1 subunits is the human angiotensin converting enzyme 2 (hACE2) (5, 8, 9). The crystallographic structures of various complexes involving the receptor binding domain (RBD) of SARS-CoV strains from different host species and the ACE2 receptor from various animal species have been described earlier (1, 10–13). Recently, the cryo-EM structure of the SARS-CoV-2 spike trimer (14) and crystallographic structures of the complex between hACE2 and the SARS-CoV-2 RBD (15, 16) have been determined.

The above structural studies and additional biochemical experiments have provided insights on the similarities and differences of the two complexes. The SARS-CoV and SARS-CoV-2 RBDs have similar folds, with an RMSD of 1.2 Å between C_*α*_ atoms (15, 16). Overall, the RBD sequences differ at 48 positions, with 34 differences located in the RBMs. Eight preserved residues and six residues with similar physicochemical properties are located in structurally equivalent positions and form similar contacts with hACE2 (Figure 1). CoV-2/CoV positions L455/Y442, F486/L472, Q493/N479, S494/D480 and N501/T487 affect the recognition of the ACE2 receptor and the range of hosts infected by the SARS-CoV virus (12, 16). Residues L455/Y442, S494/D480 and Q493/N479 are located near key salt bridge K31-E35 (“hotspot 31”) on the hACE receptor. N501/T487 interact with a second key hACE2 salt bridge D38-K353 (“hotspot 353”). The remaining differences are at positions not in direct contact with hACE2. A large number contain charge modifications, which perturb long-range interactions with the negatively charged hACE2, and might contribute to the relative affinity of the complexes.

**Figure 1:**
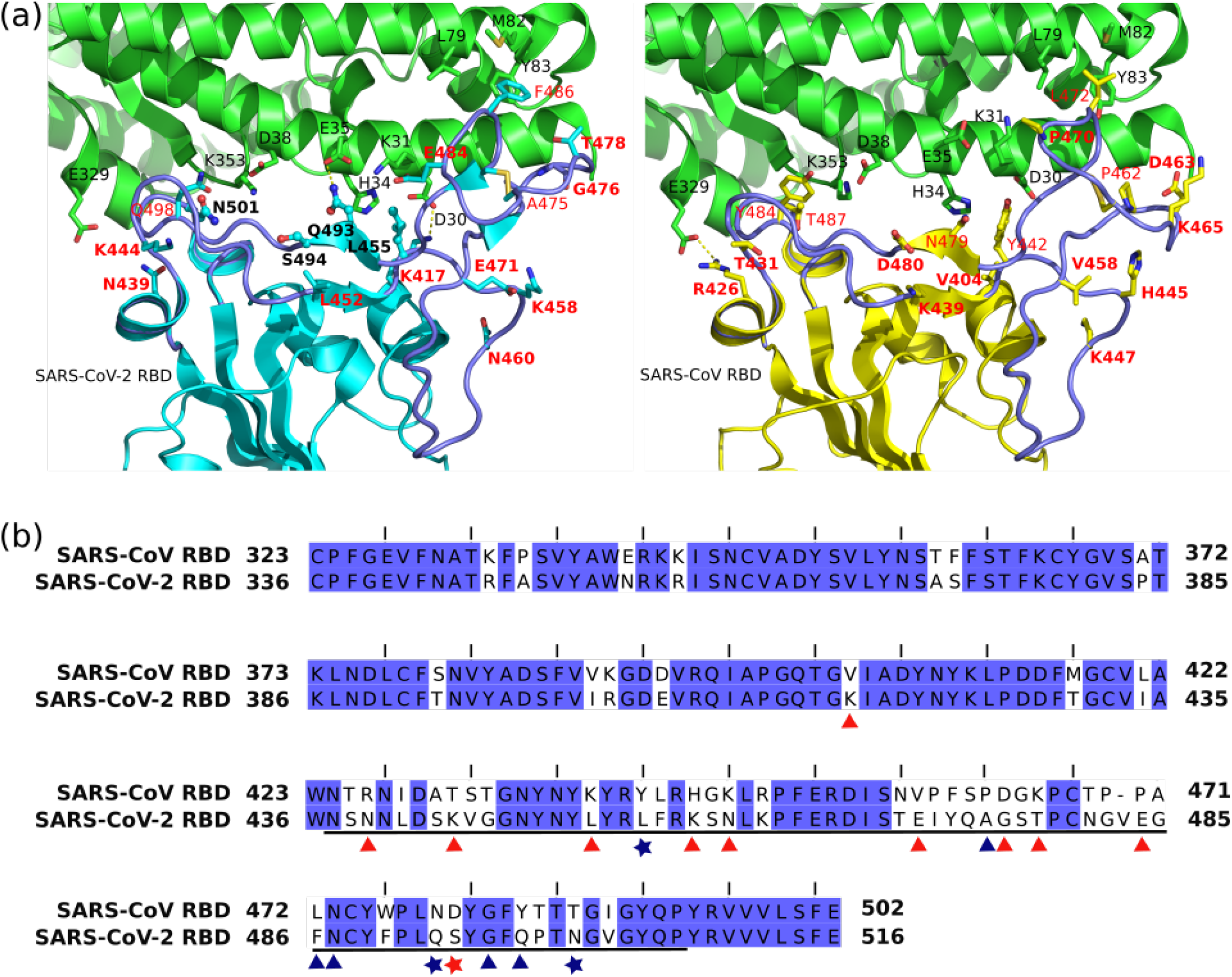
Crystallographic structures and sequence alignment of SARS-CoV and SARS-CoV-2 RBDs. (a) Side-by-side view of crystallographic structures of the SARS-CoV-2 RBD (cyan) and SARS-CoV RBD (yellow) in complex with human ACE2 (green) (8, 15). RBMs are shown in purple tube representation. ACE2 residues are indicated by black labels; positions with substitutions between CoV and CoV-2 discussed in the text are in red labels (charge-modifying substitutions in bold red); CoV-2 residues targeted by our design are shown in stick-and-ball representation, with bold black labels. (b) Sequence alignment of the SARS-CoV and SARS-CoV-2 RBDs. The RBM is underlined. Residues with substitutions discussed in the text are indicated by triangles; charge substitutions are indicated by red triangles; residues employed in design are indicated by stars.

The stronger affinity of the SARS-CoV-2 S1 protein for hACE2 may be linked to the higher human infectivity of the SARS-CoV-2 virus (15, 16). The differences between the two RBMs contribute to changes in the affinities of the two complexes. It is thus important to identify residues that contribute to the observed differences in affinity of the SARS-CoV-2 and SARS-CoV RBDs for hACE2, and search for mutations that might enhance the SARS-CoV-2 affinity.

In the present work, we study complexes of the hACE2 receptor with the RBDs of the human SARS-CoV-2 and human SARS-CoV viruses. In what follows, we refer to these complexes as “the CoV-2 complex” and “the CoV complex”, respectively. Using all-atom molecular dynamics (MD) simulations, we evaluate the contributions of the various RBD and hACE2 residues to the affinities of the two complexes. Several charge-modifying substitutions at remote positions in the CoV-2 RBD tend to stabilize or destabilize the CoV-2 complex relative to the CoV complex, but their combined effect is small; thus, the relative affinity is mainly determined by mutations at positions, which contact hACE2 in the CoV-2 complex.

We also perform exhaustive computational sequence design (CPD) calculations at key positions 455, 493, 494 and 501 of the SARS-CoV-2 RBM. The CPD calculations employ a physics-based energy function and a recent methodology for the flattening of the sequence space landscape (17–19) that is closely analogous to the Wang-Landau approach (20). The top affinity sequences contain a glutamic or aspartic acid at position 493, in place of glutamine, and nonpolar, aromatic or polar (threonine) residues at position 494, in place of serine. Substitutions at positions 455 and 501 have a smaller effect on affinity; thus CoV-2 residues L455 and N501 seem to be already optimal choices. Some of the observed substitutions are see in virus sequences encountered in bat and pangolin. Our results might be used to identify potential virus strains with increased hACE2 affinity, or peptides and peptidomimetic compounds with the potential to inhibit the CoV-2 S1:hACE2 complex.

## METHODS

### Molecular Dynamics Simulations

#### Simulation systems

We investigated complexes of human ACE2 with the SARS-CoV or SARS-CoV-2 RBDs. The crystal structures (PDB ID 2AJF (16) and 6M0J (15)) contain the hACE2 segment 323 - 502, the SARS-CoV2 segment 333-526, the SARS-CoV segment 323 - 502, Asn-linked glycans (N-Acetyl-D-Glucosamine molecules) at positions 53 (SARS-CoV complex), 90, 322, 546 in hACE2, position 330 in SARS CoV, position 343 in SARS-CoV-2, two metal ions Zn^++^ and Cl^−^, and numerous water molecules. The structures were truncated by deleting hACE2 residues or SARS-CoV-2(CoV) residues located more than 25 Å from SARS-CoV-2(CoV) or hACE2, respectively. The resulting complexes contained hACE2 segments 19-109, 293-421, 547-565, SARS-CoV segments 326-343, 353-368, 384-499 (RBM) and SARS-CoV-2 segments 339-356, 366-381, 397-513 (RBM) and had 6250-6251 protein atoms. The zinc ions, all glycans and crystallographic waters within 4 Å from the complexes were retained. The disulphide bridges C344-C361 in hACE2, C366-C419, C467-C474 in SARS-CoV, and C379-C432, C480-C488 in SARS-CoV-2 were also retained. Acetylated and N-methylated blocking groups were respectively attached at the N- and C-terminal ends of the proteins. Titratable residues were assigned their most common ionization state at physiological pH, with the exception of hACE2 residue Asp382, which was protonated. In both crystallographic complexes, residues Asp382 and Asp350 are in direct contact (C_β_-C_β_ ~ 4 Å) and their interactions with nearby residues and a crystallographic water (~ 3.5 Å) suggest that one of them is protonated. Calculations with the empirical model Propka (21) predicted highly elevated pKa values (8.8-9.7 for Asp382 and 6.6-6.0 for Asp350), suggesting that Asp382 is protonated.

The two complexes were solvated in a truncated octahedral box of explicit water molecules; the hydration layer around each complex had a minimal width of 13 Å. Potassium ions were included to neutralize the simulation systems. Initial hydrogen atom coordinates were determined by the HBUILD facility (22). All system setups were performed with the CHARMM-GUI interface (23).

#### Equilibration and production simulations

The solvated systems were first subjected to 800 steps of conjugate-gradient minimization. Protein and glycan heavy atoms and the Zn ion were harmonically restrained, with restraint constants gradually varied from 10 to 2 kcal/mol/Å^2^; this was followed by 200 adopted-basis Newton-Raphson minimization steps, with 1.0 kcal/mol/Å^2^ restraint constants on backbone heavy atoms and 0.1 kcal/mol/Å^2^ on sidechain heavy atoms. Additional dihedral restraints ensured that glycans kept their sugar conformation.

Following minimization, each system was equilibrated by an 1-ns simulation in the NPT ensemble at 300 K, with harmonic restraints applied on backbone and side chain heavy atoms. The simulation was divided into six segments with lengths of 3×100 ps, 2×150 ps and 400 ps, in which the harmonic restraints were gradually reduced from 10 kcal/mol/Å^2^ on all protein heavy atoms to 0.5 kcal/mol/Å^2^ on main chain atoms. In the last segment the time step was increased from 1.0-fs to 1.5-fs to 2.0-fs.

The production simulations were performed in the NPT ensemble with the NAMD program (24). All protein heavy atoms located more than 15 Å from the hACE2 or SARS-CoV-2/CoV segments were retained near their initial positions by harmonic restraints. The restraint constant was set to 0.4 kcal/mol/Å^2^ to reproduce vibrational thermal motions, as reflected by the crystallographic B-factors (65 Å^2^).

The system temperature was controlled by Langevin dynamics at 300 K, with a friction coefficient of 5 ps^−1^. The pressure was kept constant at 1 atm using a Nose-Hoover Langevin piston with a period of 200 ps (25, 26). Atomic interactions were modeled by the CHARMM36 energy function (27). The water solvent was represented by the TIP3P model (28–30). Long-range electrostatic interactions were described (every 2 steps) by the Particle Mesh Ewald method (31) using the r-RESPA multiple time-step method (32) and a general cutoff distance at 12 Å for all non-bonded interactions. We performed 3 production runs from different starting structures. Each run had a time-step of 2 fs and a total duration of 41 ns; post-processing analysis was performed with 4000 snapshots, extracted at 10-ps intervals from the last 40 ns of each trajectory.

#### Estimation of binding affinities

The total effective energies were expressed as the sum of molecular mechanics (MM) and solvation (solv) terms:

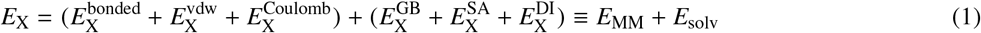

where X is the system under consideration (the complex C or the dissociated proteins P1 and P2). The MM energy terms correspond to bonded, van der Waals (vdw) and Coulombic interactions. The continuum electrostatic generalized Born (GB) term *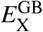* models the interaction of the solute charges with the solvent polarization. The term *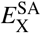* takes into account the tendency of solute atoms to be exposed or hidden from solvent via a surface area (SA) term. The term *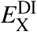* incorporates solute-water attractive dispersion interactions (33–36).

To compute these terms, we extracted 12000 coordinate frames at 10-ps intervals and removed all waters and ions. We employed the “single-trajectory” approximation, according to which the two proteins have identical conformations in the complex and unbound states. The binding affinities are

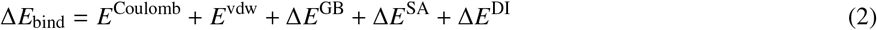

with *E*^Coulomb^ and *E*^vdw^ the intermolecular Coulomb and vdw energies. Bonded and intramolecular vdw and Coulomb energies are identical in the complex and unbound proteins and do not appear in Eq. 2. The binding affinities do not take into account changes in the translational, rotational and vibrational/conformational entropies of the two proteins upon association. These terms are expected to cancel to a large extent in the relative affinities for molecules of comparable sizes, such as the CoV-2 and CoV RBDs (37, 38).

Electrostatic intermolecular residue energies were computed by the equation

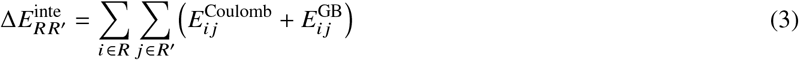

with 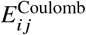 and 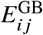 the Coulomb and GB interaction energies between atoms *i* and *j*. In our case, *R* was a CoV-2 RBM residue and *R*, was the entire hACE2, or vice-versa.

### Optimization of the CoV-2 RBM affinity for hACE2

#### Design positions

We designed key positions 455, 493, 494 and 501 in the CoV-2 RBM, to improve affinity for hACE2. The design calculations were conducted with the program Proteus (19, 39).

#### Design scaffold

The system used in the design resulted from the all-atom simulation CoV-2 complex, after removing glycans, metal ions and water molecules. The resulting complex contained 239 hACE2 residues and 157 SARS-CoV-2 RBD residues.

The initial atomic coordinates were taken from the crystallographic structure (15) and were subjected to 1000 steps of minimization. During design, positions 455, 493 and 494 were allowed to sample 18 chemical types (A, I, L, V, M, K, R, D, E, N, Q, C, S, T, F, Y, W, H(N_δ_)). Position 501 sampled 14 types; bulky sidechains F, Y, W, H were excluded, due to steric repulsions. These selections resulted in 18^3^ × 14 = 81648 possible sequences. All chemical types sampled sidechain conformations from a discrete rotamer library (40), augmented to include orientations seen in the PDB structure of the CoV-2 complex (15). 53 SARS-CoV-2 and 108 hACE2 side chains (excluding glycine and proline residues) within 15 Å of hACE2 retained the SARS-CoV-2 chemical type and sampled conformations from the same library. The remaining atoms (the entire protein backbone, including the N- and C-terminal blocking groups, cysteine residues in disulphide bridges, all glycines and prolines and all other sidechains farther than 15 Å from the SARS-CoV-2/ACE2 interface) were kept fixed.

#### Interaction Energy Matrix

The interaction energies of all side chain-backbone and sidechain-sidechain pairs, for all possible side-chain chemical types and rotamers, were pre-computed and stored in an Interaction Energy Matrix. The energy function was given from Eq. 1. The GB term corresponded to the Hawkins-Cramer-Truhlar Generalized Born (GB/HCT) approximation (41–43). It was rendered pair-wise decomposable via the Native Environment Approximation (NEA), that is described in (44–46). According to this approximation, the atomic solvation coefficients b_i_ of each residue are precalculated, with the rest of the system kept at the native sequence and conformation. The water and protein dielectric constants were set to 80.0 and 6.8; the atomic surface coefficients of non-polar, polar, ionic and aromatic atoms were set to σ_alk_ = −5 cal/mol, σ_pol_ =−8 cal/mol, σ_ion_ =−9 cal/mol, σ_aro_ =−12 cal/mol (33).

#### Adaptive flattening of the SARS-CoV-2 RBD apo state

We first performed a design simulation of the unbound SARS-CoV-2 RBD domain, in which we derived bias potentials that rendered all allowed chemical types at positions 455, 493, 494 and 501 equiprobable. The procedure is analogous to the Wang-Landau approach (20) and has been described in detail in refs (17–19). At a particular simulation time *t*, the bias potentials have the form

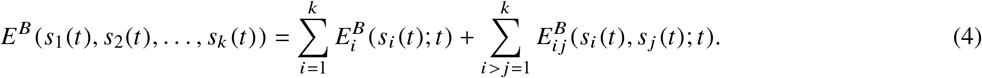

The above sums are over the four mutable positions; *s_i_* (*t*) is the sidechain type at position *i* at simulation time *t*; 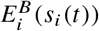 is the biasing potential for sidechain type *s_i_* at position *i*, and 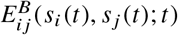 is the biasing potential for pairs of side-chain types *s_i_* and *s _j_* at positions *i* and *j*, at simulation time *t*.

The bias potentials are updated at regular time intervals *T*. During an update at simulation time *t* = *nT*, the sequence *s*_1_ (*nT*),…,*s_k_* (*nT*) is penalized by adding the following energy increments to the corresponding bias potentials:

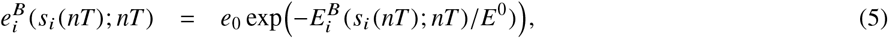

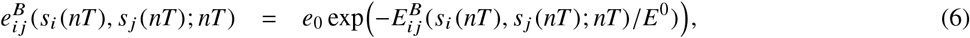

where *e*_0_ and *E*^0^ are constants with the dimension of energy. As the simulation proceeds, the biasing potentials grow (with exponentially decreasing increments), reducing the probabilities of frequently appearing sequences and flattening the sequence space. The resulting total bias potentials at simulation time t are

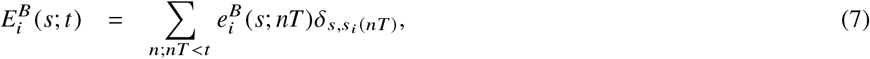

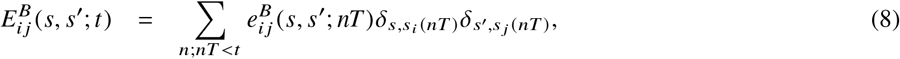

where *δ*_*a,b*_ is Kronecker’s delta.

In the current work, the bias potentials were updated every 1000 MC steps. The parameters *e*_0_ and *E*_0_ were respectively set to 0.2 kcal/mol and 100 kcal/mol for single-position biases and 0.1 kcal/mol, 40 kcal/mol for two-position biases. The biases were updated via 3 × 10^8^ MC steps of the free SARS-CoV-2 at a temperature of 300 K, using single- and double-position moves.

#### Biased simulations of the ACE2 complex

The resulting biased potentials are approximately equal to the negative folding energies of the apo (A) SARS-CoV-2 RBD state (18). Using the same biasing potentials, a second design simulation is conducted for the CoV-2 complex (C). Since the biasing potentials subtract from the free energies of the complex the unbound-protein free energies, the design promotes the selection of sequences with good binding affinities. The populations *p_C_* (*S*) and *p_A_* (*S*) of a sequence *S* in the simulations of the complex and the apo states are used to compute the binding free energy of a sequence *S* relative to a reference sequence *S*_ref_:

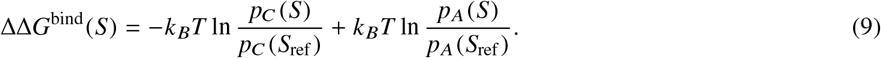

The probabilities entering Eq. 9 are biased. However, if the same biasing potentials are used in the design simulations of the apo state and the complex (as done here), the biasing terms cancel out in the binding free energies (18). Thus, Eq. 9 yields the unbiased binding free energy of sequence S relative to a reference sequence *S*_ref_ (e.g., the native SARS-CoV-2 sequence).

#### Details of MC simulations

The biasing MC simulations of the apo SARS-CoV-2 RBD and the CoV-2 complex had a length of 10^9^ Replica Exchange Monte Carlo steps. We employed four replicas at kT = 0.6 kcal/mol, 0.9 kcal/mol, 1.3 kcal/mol and 1.8 kcal/mol; swaps between neighboring replicas were attempted every 2000 MC steps. The simulation started from a randomly selected sequence/conformation state using the random number generator routine mt19937 from GSL. Side chains more than 7 Å from any atom of the 4 mutable positions were retained to their initial conformations. At each MC step, a chemical type/rotamer modification was randomly chosen at one or two positions. The associated energy difference was computed by the IEM, and the modification was accepted or rejected according to the Metropolis criterion (47). The frequencies of attempted changes were as follows: one-position rotamer changes 57% and chemical-type changes 11%, two-position rotamer/rotamer changes 23%, type/rotamer changes 6% and type/type changes 3%. Residue pairs with interaction energies smaller in absolute value than 3 kcal/mol were not considered for two-position changes. At the end of the simulation, sequences were collected from the 0.6 kcal/mol replica for analysis.

## RESULTS

### All-atom MD simulations of the CoV-2 and CoV complexes

#### Description of the crystallographic CoV-2 and CoV complexes

Figure 1(a) displays the crystallographic structures of the two complexes. The CoV-2 RBD spans the S1 protein region 333-526 and is folded into an antiparallel, five-strand β-sheet (strands β1, β2, β3, β4, β7), connected via three short helices (α1, α2, α3) and loops; recognition of the hACE2 receptor is achieved via region 438-506 (the receptor binding motif or RBM), that is inserted between strands β4 and β7 and consists of two short strands (β5 and β6), two helices (α4, α5) and loops (15, 16, 48). In SARS-CoV, the corresponding RBD extends in the region T425-S492 and the RBM in the region T425-Q492.

A sequence alignment of the CoV-2 and SARS-CoV RBDs is included in Figure 1(b); aligned residues occupy structurally equivalent positions in the two RBDs. Using a cut-off distance of 4 Å, 17 SARS-CoV-2 residues and 16 SARS-CoV residues in the two RBMs form crystallographic intermolecular contacts with hACE2 (8, 15, 16). Eight CoV-2/CoV residues are identical (Y449/Y436, Y453/Y440, N487/473, Y489/Y475, G496/G482, T500/T486, G502/G488, Y505/Y491) and six residues are modified, but retain similar biochemical properties (L455/Y442, F456/L443, F486/L472, Q493/N479, Q498/Y484, N501/T487). These residues form similar hydrogen-bonding or nonpolar contacts with hACE2 (Table 1). Hotspot-31 residue K31 contacts the structurally equivalent residues Y489/Y475. At hotspot 353, residue D38 forms a hydrogen bond with the structurally equivalent residues Y449/Y436; K353 forms a main chain hydrogen bond with residues G502/G488 and nonpolar contacts with Y505/Y491.

**Table 1:** Common crystallographic intermolecular hydrogen-bonds and contacts in the CoV-2 and CoV complexes.

| CoV-2<br>residue | CoV<br>residue | hACE2<br>residue | h-bond<br>CoV-2/CoV | contact<br>CoV-2/CoV |
| --- | --- | --- | --- | --- |
| Y449 | Y436 | D38 | 1 sc-sc<sup>a</sup> |  |
| Y449 | Y436 | Q42 | sc-sc |  |
| Y453 | Y440 | H34 |  | p-np<sup>a</sup> |
| F456 | L443 | T27 |  | np-np<sup>a</sup> |
| F486 | L472 | L79 |  | np-np |
| N487 | N473 | Q24 | sc-sc |  |
| N487 | N473 | Y83 | sc-sc |  |
| Y489 | Y475 | T27 |  | np-np |
| Y489 | Y475 | F28 |  | p-np |
| Y489 | Y475 | K31 |  | np-np |
| Y489 | Y475 | Y83 | sc-sc |  |
| G496 | G482 | K353 |  | p-np |
| Q498 | Y484 | Y41 |  | np-np |
| Q498 | Y484 | Q42 |  | p-p |
| T500 | T486 | Y41 | sc-sc |  |
| T500 | T486 | N330 |  | np-np |
| T500 | T486 | D355 |  | p-p |
| T500 | T486 | R357 |  | np-p |
| **N501** | T487 | Y41 | mc-sc |  |
| G502 | G488 | K353 | sc-mc |  |
| G502 | G488 | G354 |  | np-p |
| Y505 | Y491 | E37 | sc-sc |  |
| Y505 | Y491 | K353 |  | np-np |
| Y505 | Y491 | G354 |  | np-p |

Structurally equivalent residues in the CoV-2 and CoV RBMs are included in the same row. Designed positions are in bold.
<sup>a</sup> “sc” and “mc” denote hydrogen-bonds involving side chain and mainchain atoms, respectively; “np” and “p” denote contacts involving nonpolar and polar moieties.

Contacts observed only in one of the two complexes are collected in Table 2. In the CoV-2 complex, hACE2 residue D30 forms a stabilizing salt-bridge with K417 (outside the RBM) and an extra hydrophobic contact with F456; at hotspot 31, hACE2 K31 makes a polar-nonpolar contact with Y442, H34 a nonpolar contact with residue L455 and E35 a hydrogen bond with Q493; at hotspot 353, residue K353 makes a hydrogen bond with G496. Residues G446 and Y505 form new intermolecular hydrogen bonds, respectively, with Q42 and R393; A475 makes a contact with Q24; F486 with M82 and Y83. In the CoV complex, K353 forms a nonpolar contact with T487; R426 forms two hydrogen bonds with E329 and Q325 and T486 forms a hydrogen bond with N330; Y484 and T487 make nonpolar contacts with L45 and Y41; G488 and I489 make nonpolar/polar contacts with D355 and Q325.

**Table 2:** Distinct crystallographic hydrogen-bonds and contacts in the CoV-2 and CoV complexes.

| CoV-2<br>residue | CoV<br>residue | hACE2<br>residue | h-bond<br>CoV-2/CoV | contact<br>CoV-2/CoV |
| --- | --- | --- | --- | --- |
| K417 | V404 | D30 | sc-sc/-<sup>a,b</sup> |  |
| N439 | R426 | Q325 | -/sc-sc |  |
| N439 | R426 | E329 | -/sc-sc |  |
| G446 | T433 | Q42 | mc-sc/-<sup>a,b</sup> |  |
| **L455** | Y442 | K31 |  | -/p-np<sup>a,b</sup> |
| **L455** | Y442 | H34 |  | np-np/-<sup>a,b</sup> |
| F456 | L443 | D30 |  | np-np/- |
| A475 | P462 | Q24 |  | p-np/- |
| F486 | L472 | M82 |  | np-np/np-p |
| F486 | L472 | Y83 |  | np-np/- |
| **Q493** | N479 | K31 |  | p-np/- |
| **Q493** | N479 | E35 | sc-sc/- |  |
| G496 | G482 | K353 | mc-sc/- |  |
| Q498 | Y484 | L45 |  | -/np-np |
| T500 | T486 | L45 |  | -/p-np |
| T500 | T486 | N330 | -/mc-sc |  |
| **N501** | T487 | K353 |  | p-np/np-np |
| G502 | G488 | D355 |  | -/p-p |
| V503 | I489 | Q325 |  | -/np-p |
| Y505 | Y491 | R393 | sc-sc/- |  |

Structurally equivalent residues in the CoV-2 and CoV RBMs are included in the same row. Designed positions are in bold.
<sup>a</sup> “sc” and “mc” denote hydrogen-bonds involving side chain and mainchain atoms, respectively; “np” and “p” denote contacts involving nonpolar and polar moieties.
<sup>b</sup> The absence of a hydrogen-bond or contact in a complex is denoted by “-”.

The RBD sequences contain also charge-modifying substitutions at positions not in direct contact with hACE2 in the complexes. CoV-2/CoV substitutions K444/T431, K458/H445, G476/D463 and S994/D480 introduce a positive charge or eliminate a negative charge in the COV-2 RBM; substitutions N439/R426, L452/K439, N460/K447, E471/V458, T478/K465, E484/P471 eliminate a positive charge or insert a negative charge (Figure 1). As discussed below, residues at these positions form longer-range polar interactions with hACE2 and contribute to the affinities of the two complexes.

#### Interactions in the MD simulations of the CoV-2 and CoV complexes

The intermolecular crystallographic contacts of Tables 1-2 were maintained in the simulations, with the exception of CoV-2 contact F456-D30 that was replaced by L455-D30. Some new intermolecular contacts were also observed. CoV-2/CoV residues A475/P462 formed contacts with Q24 and T27; residues G476/P462 formed a new contact with S19. An extended hydrophobic packing was observed near hotspot 353, involving T500/T486, N501/T487, G502/G488, Y505/Y491 and ACE2 residues N330, K353, G354, D355. New intermolecular contacts only in the CoV-2 complex involved residue pairs G476-Q24 and G496-D38; in the CoV complex, new contacts involved pairs N473-Q24 and G488-T324. Overall, the CoV-2 RBD formed 21 non-polar contacts with hACE2, compared to 20 contacts in the CoV complex. The two hotspots (intermolecular hACE2 salt bridges K31-E35 and D38-K353) were very stable, with occupancies in the range 100%-85%.

Statistics of intermolecular hydrogen-bonds are collected in Table 3. Residues N487/N473 form two hydrogen bonds with Y83 and Q24, residues Y489/Y475 form a second hydrogen bond with Y83, G502/G488 a hydrogen bond with K353, and T500/T486 with Y41. In general, common hydrogen bonds are more stable in the CoV-2 simulations. The intermolecular salt bridge K417-D30 (54.5% occupancy) and the hydrogen bonds K31-Q493 (27.6%), E35-Q493 (51.5%), K353-G496 (66.4%) and K353-Q498 (34.6%) are only observed in the CoV-2 complex. In the CoV complex, hotspot-31 residues do not form intermolecular hydrogen-bonds; K353 forms a single hydrogen bond with G488 (45.2% occupancy) and D38 forms three infrequent hydrogen bonds with residues N479, Y436 and Y484 (occupancies 29.3%-15.4%).

**Table 3:**
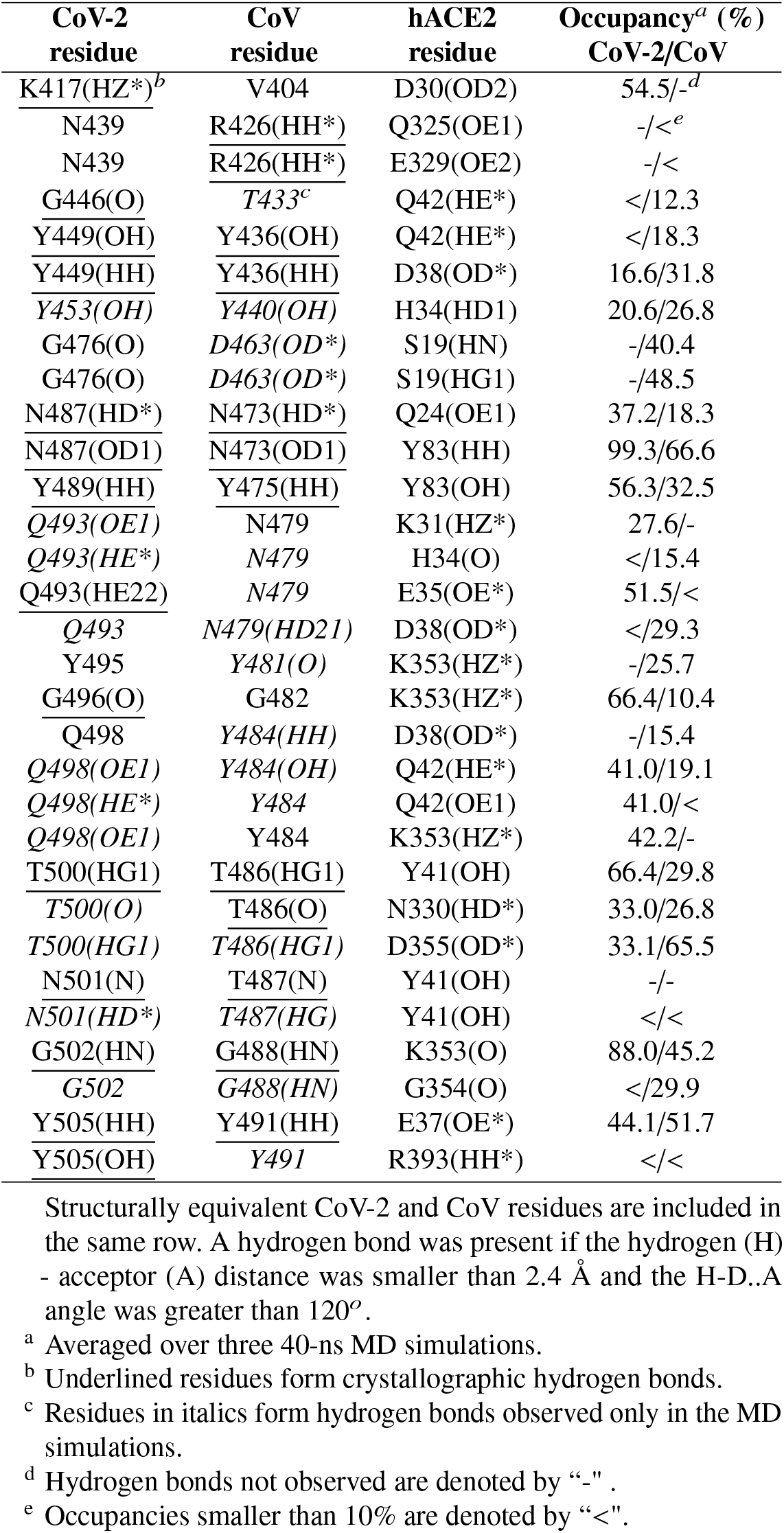
Intermolecular hydrogen bonds in the CoV-2 and CoV complexes.

| CoV-2<br>residue | CoV<br>residue | hACE2<br>residue | Occupancy<sup>a</sup> (%)<br>CoV-2/CoV |
| --- | --- | --- | --- |
| K417(HZ*)<sup>b</sup> | V404 | D30(OD2) | 54.5/-<sup>d</sup> |
| N439 | R426(HH*) | Q325(OE1) | -/<<sup>e</sup> |
| N439 | R426(HH*) | E329(OE2) | -/< |
| G446(O) | T433<sup>c</sup> | Q42(HE*) | </12.3 |
| Y449(OH) | Y436(OH) | Q42(HE*) | </18.3 |
| Y449(HH) | Y436(HH) | D38(OD*) | 16.6/31.8 |
| Y453(OH) | Y440(OH) | H34(HD1) | 20.6/26.8 |
| G476(O) | D463(OD*) | S19(HN) | -/40.4 |
| G476(O) | D463(OD*) | S19(HG1) | -/48.5 |
| N487(HD*) | N473(HD*) | Q24(OE1) | 37.2/18.3 |
| N487(OD1) | N473(OD1) | Y83(HH) | 99.3/66.6 |
| Y489(HH) | Y475(HH) | Y83(OH) | 56.3/32.5 |
| Q493(OE1) | N479 | K31(HZ*) | 27.6/- |
| Q493(HE*) | N479 | H34(O) | </15.4 |
| Q493(HE22) | N479 | E35(OE*) | 51.5/< |
| Q493 | N479(HD21) | D38(OD*) | </29.3 |
| Y495 | Y481(O) | K353(HZ*) | -/25.7 |
| G496(O) | G482 | K353(HZ*) | 66.4/10.4 |
| Q498 | Y484(HH) | D38(OD*) | -/15.4 |
| Q498(OE1) | Y484(OH) | Q42(HE*) | 41.0/19.1 |
| Q498(HE*) | Y484 | Q42(OE1) | 41.0/< |
| Q498(OE1) | Y484 | K353(HZ*) | 42.2/- |
| T500(HG1) | T486(HG1) | Y41(OH) | 66.4/29.8 |
| T500(O) | T486(O) | N330(HD*) | 33.0/26.8 |
| T500(HG1) | T486(HG1) | D355(OD*) | 33.1/65.5 |
| N501(N) | T487(N) | Y41(OH) | -/- |
| N501(HD*) | T487(HG) | Y41(OH) | </< |
| G502(HN) | G488(HN) | K353(O) | 88.0/45.2 |
| G502 | G488(HN) | G354(O) | </29.9 |
| Y505(HH) | Y491(HH) | E37(OE*) | 44.1/51.7 |
| Y505(OH) | Y491 | R393(HH*) | </< |

Structurally equivalent CoV-2 and CoV residues are included in the same row. A hydrogen bond was present if the hydrogen (H) - acceptor (A) distance was smaller than 2.4 Å and the H-D..A angle was greater than 120°.
<sup>a</sup> Averaged over three 40-ns MD simulations.
<sup>b</sup> Underlined residues form crystallographic hydrogen bonds.
<sup>c</sup> Residues in italics form hydrogen bonds observed only in the MD simulations.
<sup>d</sup> Hydrogen bonds not observed are denoted by "-".
<sup>e</sup> Occupancies smaller than 10% are denoted by "<".

#### Residue interaction energy analysis of the CoV-2 and CoV complexes

Figure 2 displays difference values for selected residue intermolecular energies in the CoV-2 and CoV complexes. Negative values indicate stronger intermolecular interactions in the CoV-2 complex. Values at non-conserved positions show whether a specific substitution improves or weakens intermolecular interactions. The energies are averaged over three independent 40-ns MD trajectories of each complex, with the standard deviation of the means in error bars.

**Figure 2:**
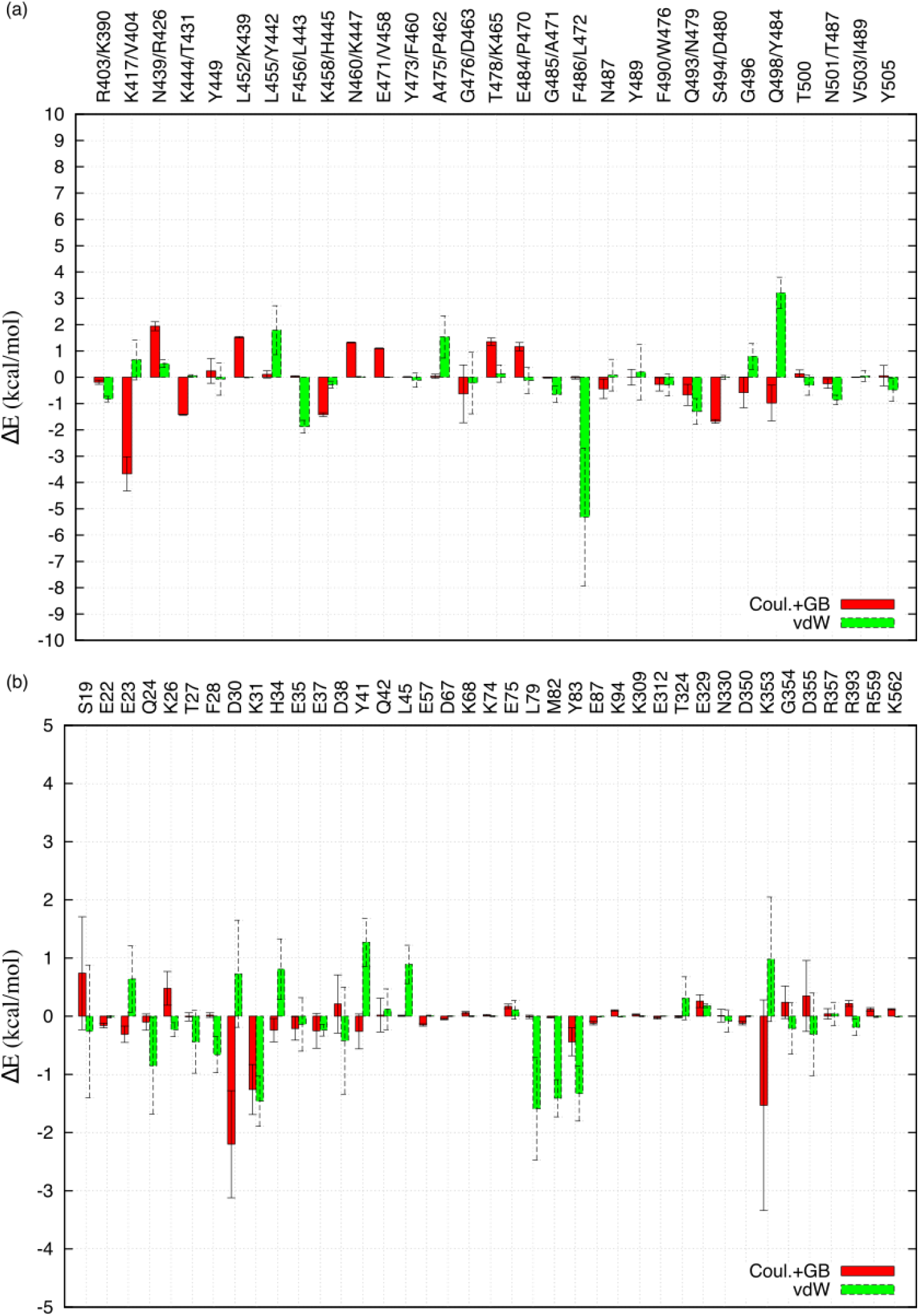
Residue intermolecular interaction energy difference profiles. Negative values signify stronger interactions in the CoV-2 complex. (a) CoV-2/CoV RBD residue profile; (b) hACE2 residue profile.

The CoV-2/CoV residue profiles are shown in Figure 2(a). We first discuss the differences in the electrostatic (Coulomb+GB) intermolecular energies. Substitutions K417/V403, K444/T431, K458/H445, G476/D463, S994/D480 increase the positive-charge of the CoV-2 RBD and strengthen interactions with the excess negative charge of the hACE2 receptor (11 positively and 14 negatively charged hACE2 residues are located within 15 Å of the CoV-2 RBM). Substitutions N439/R426, L452/K439, N460/K447, E471/V458, T478/K465, E484/P471 reduce the positive charge and weaken interactions with hACE2. These residues are shown in Figure 1(a). With the exception of K417 (forming a salt-bridge with CoV-2 S30), R426 (hydrogen-bonded to Q325 and E329 in CoV) and D463 (interacting with CoV S19), other positions do not contact hACE2. Due to cancellations among the above residue components, their total contribution to the relative affinity is small (−0.4 kcal/mol). Additional negative contributions are due to charge-preserving substitutions Q493/N479 (−0.7 kcal/mol), Q498/Y484 (−1.0 kcal/mol), and conserved residues N487 (−0.4 kcal/mol), G496 (−0.6 kcal/mol). All these residues form more stable hydrogen bonds with hACE2 in the CoV-2 simulations (Table 3).

With respect to vdw interactions, the most negative contribution is associated with substitution F486/L472, due to improved contacts of F486 with hACE2 residues L79, M82 and Y83. The most positive value is due to substitution Q498/Y484; in the CoV simulations, Y484 forms better intermolecular interactions with Y41.

The above analysis shows that charge-modifying substitutions between the CoV and CoV-2 RBDs do not always favor the CoV-2 complex, but tend to cancel each other and contribute little to the relative affinity of the CoV-2 complex. Some substitutions may be chosen for stability reasons, as they eliminate or create pairs of opposite charges at or near salt-bridge distances (Figure 1(a)): the CoV pairs D463-K465, K439-D480 and H445-V458 are transformed to G476-T478, L452-S494 and K458-E471 in CoV-2. Overall, the observed charge balance prevents the accumulation of excessive positive charge on the SARS-CoV-2 RBM and might help the SARS-CoV-2 S1 protein to escape recognition by molecules other than the hACE2 receptor. Instead, the improved CoV-2 S1 affinity for hACE2 relies mainly on substitutions at the interface of the CoV-2 complex, notably K417/V404 and F486/L472.

The difference profiles of selected hACE2 residues are displayed in Figure 2(b). Most electrostatic components are small, suggesting that the majority of hACE2 residues do not differentiate significantly between the CoV and CoV-2 RBDs, even if they are charged. This can be attributed to the fact that the CoV-2 and CoV RBMs have similar net charges (respectively +1 and +2 within 15 Å of hACE2). Thus, maintaining a constant charge might assist the virus RBM to prevent recognition by molecules other than hACE2. The largest (in absolute value) negative electrostatic contributions are associated with D30 (hydrogen-bonds with K417), hotspot-31 K31 (hydrogen-bonds to Q493) and hotspot-353 K353 (hydrogen bonds to G496, Q498 and G502). The largest negative vdw energies are associated with K31, L79, M82 and Y83. Lysine K31 makes more favourable vdw interactions with CoV-2 residues F490 and F456. The other three residues form a hydrophobic pocket that makes improved non polar contacts with CoV-2 residue F486.

Using Eq. 2, we estimate that the CoV-2 complex has a lower total binding affinity by 4.0±2.4 kcal/mol. The corresponding experimental estimate is in the range −0.9 to −1.1 kcal/mol, based on the reported dissociation constants in (15, 16). Thus, the computational estimate ranks correctly the two complexes, but overestimates somewhat the relative CoV-2 affinity.

### Computational optimization of the SARS-CoV-2 RBM

We optimized the SARS-CoV-2 composition at RBM positions 455, 493, 494 and 501, searching for mutants with improved binding affinity for hACE2. Residues at these positions interact with the “hotspot” intramolecular salt-bridges K31-E35, K353-D38 on the hACE2 α1 helix, and have been characterized as critical for the recognition of hACE2 (15, 16). The methodology is outlined in the methods.

#### Design and filtering procedure

##### Step 1: Flattening of the energy landscape of the apo CoV-2 RBD

We first derived bias potentials that flattened the sequence landscape at positions 455, 493, 494 and 501 of of the apo SARS-CoV2 RBD. Figure 3(a) displays a logo of the sequence composition after flattening the landscape. All 18 chemical types were accepted at positions 455, 493, 494 and all 14 allowed chemical types were accepted at position 501. A total of 81643 out of 81648 possible sequences were sampled in the simulations, with 60660 sequences appearing more than 5000 times. The CoV-2 wt sequence “LQSN” was selected 15651 times, and the CoV wt sequence “YNDT” 1264 times.

**Figure 3:**
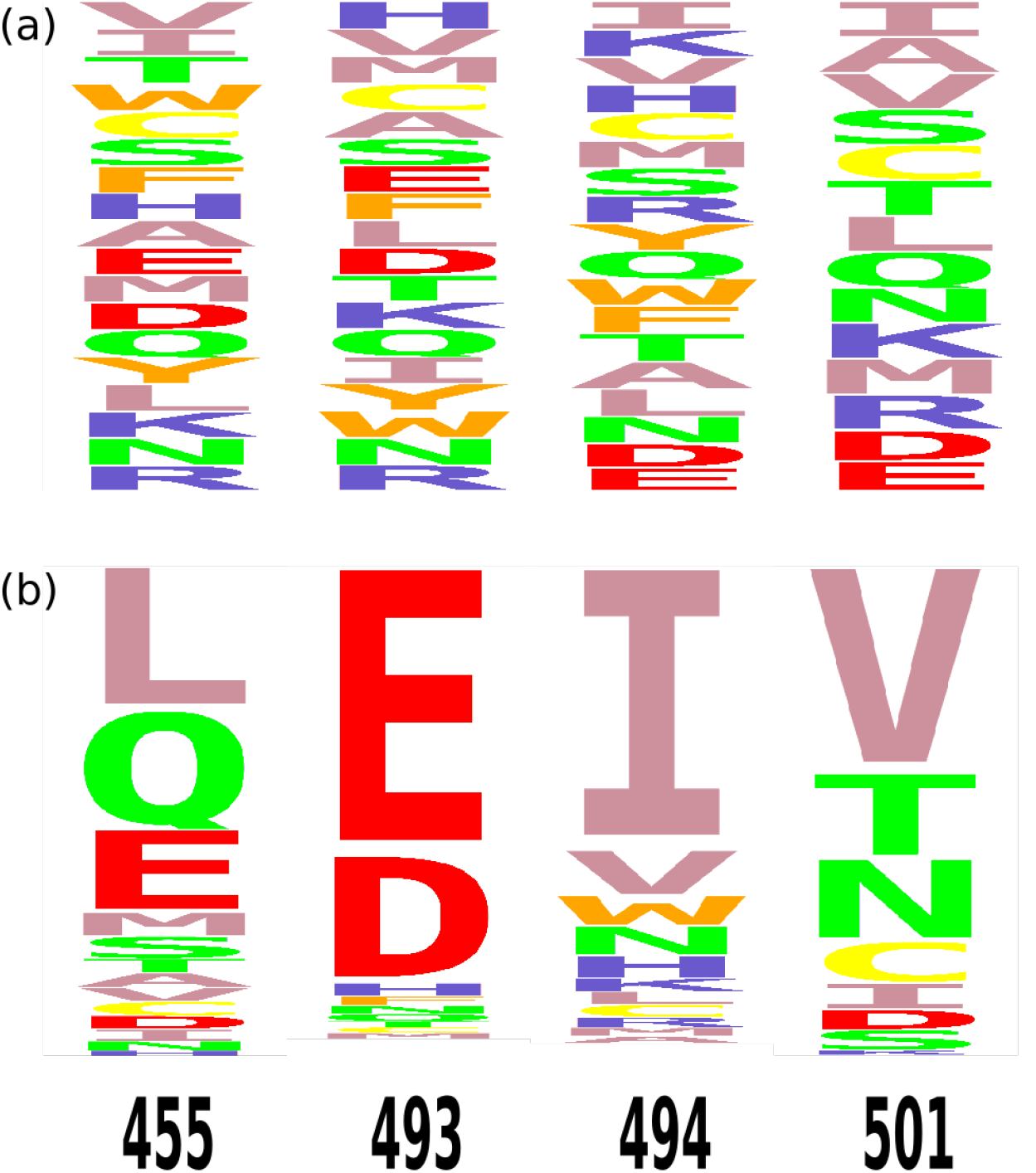
Logos of designed sequences. (a) Sequence composition of the flattened apo CoV-2 state; (b) Sequence composition of the CoV-2 complex, using the flattening biases. Large letters signify residue types encountered in sequences of better affinity for hACE2, relative to the CoV-2 sequence LQSN.

##### Step 2: Biased simulations of the apo CoV-2 RBD and the CoV-2 complex

The above bias potentials approximate the negative free energies of the apo CoV-2 RBD (18). Using these potentials, a second design simulation was conducted with the CoV-2 complex. Since the biasing potentials subtract the apo from the complex free energies, the design promotes sequences with good binding affinities. In the calculations of the CoV-2 complex, a total of 29668 out of 81648 sequences were sampled, with 6350 sequences appearing more than 5000 times. The CoV-2 sequence L455/Q493/S494/N501 (LQSN) was sampled 6347 times, whereas the CoV sequence Y455/N493/D494/T501 was not sampled. A total of 5596 common sequences were visited at least 5000 times in the simulations of the complex and apo state. Using Eq. 9, we estimated that 4704 out of the 5596 sequences had better binding affinity than the native sequence.

#### Discussion of the designed sequences

Figure 3(b) displays a logo with the composition of the 4704 sequences with improved affinities relative to the CoV-2 sequence LQSN. The resulting chemical types are Boltzmann-weighted by the sequence binding affinities; large letters signify types most frequently encountered in the highest-affinity sequences. Position 455 is most frequently occupied by a leucine (L) as in the CoV-2 RBM, a glutamine (Q) or a glutamic acid (E); position 493 is occupied by negatively charged aminoacids (D, E), position 494 by hydrophobic (I, V), aromatic (W) and polar (N) aminoacids, and position 501 by V, T (as in the CoV RBM) and N (as in the CoV-2 RBM). Figure 4(a) lists the 20 highest-affinity sequences. The top sequence is the triple mutant L455/E493/I494/V501 (LEIV).

**Figure 4:**
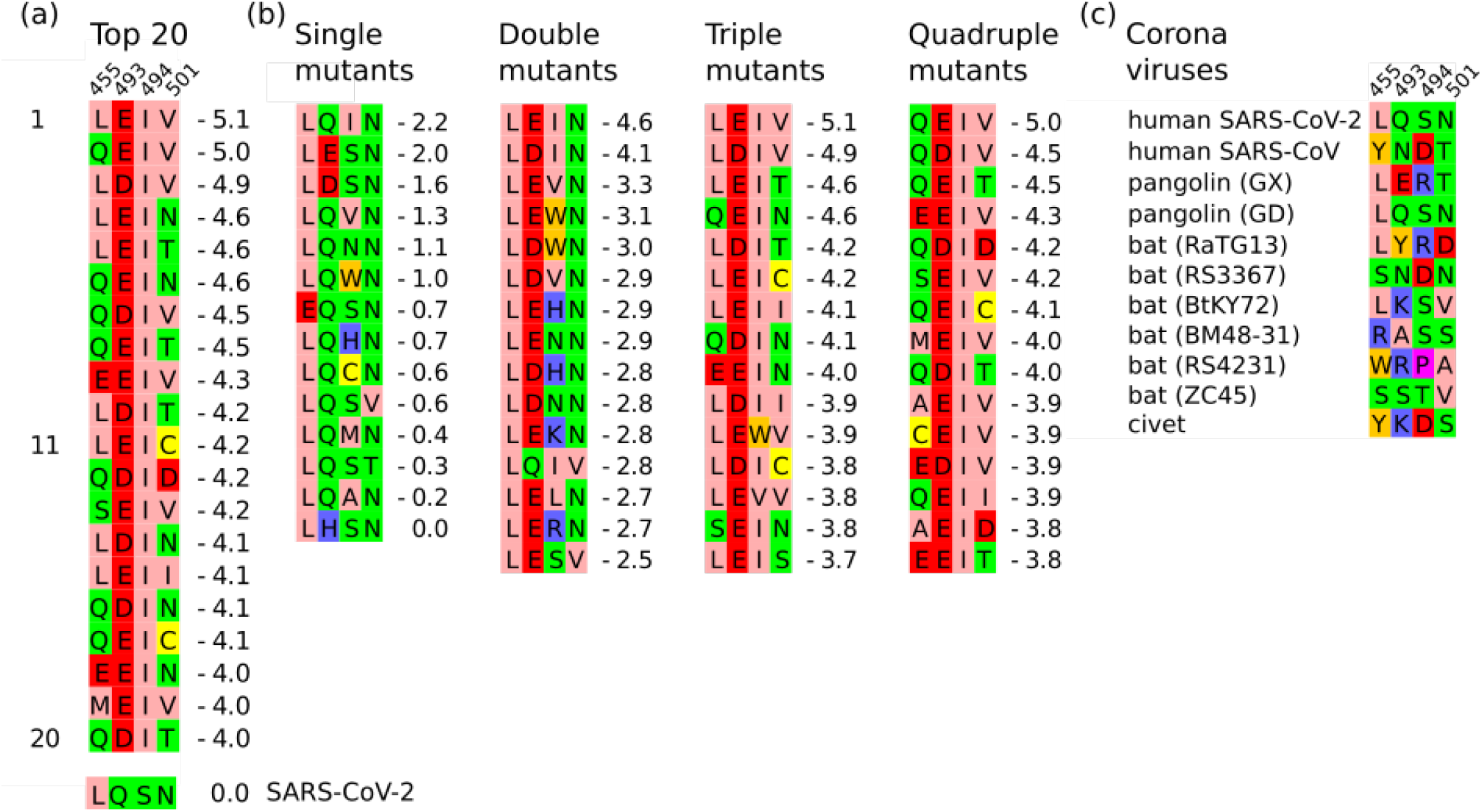
(a) Top 20 affinity sequences. (b) Top sequences, grouped by similarity with the CoV-2 sequence LQSN. (c) Sequence composition of various coronaviruses at designed positions 455, 493, 494, 501 (SARS-CoV-2 numbering) (16, 49).

Figure 4(b) displays the highest-affinity sequences, organized in terms of their similarity with the CoV-2 sequence. Single-point mutants are listed in the leftmost column. The best three sequences contain the mutations Q493E/D or S494I; each of the three mutations is predicted to improve the CoV-2 affinity for hACE2 by 1.6–2.2 kcal/mol. Position 494 displays the highest variability, with eight point mutations. Most favorable choices insert nonpolar (isoleucine, valine) or aromatic (tryptophane) sidechains, or substitute CoV-2 serine by another polar aminoacid (asparagine). At positions 455 and 501, mutations are sparse and have a weak effect on affinity. Mutations L455E and N501V improve affinity by 0.7–0.6 kcal/mol; a small improvement in affinity (0.3 kcal/mol) is also predicted for the mutation N501T, which restores at that position the chemical type encountered in the CoV RBD.

The top 15 double-mutants are displayed in the second column of Figure 4(b). Most sequences result from a combination of the best single-point mutations, i.e. contain residues E/D at position 493 and residues I, V and W at position 494. The best double mutant (LEIN) is among the four top sequences Figure 4(a), with a relative affinity of −4.6 kcal/mol.

The top triple mutants are displayed in the next-to-last column of Fig. 4(b). The top mutants combine the 493E/D and 494I residues encountered in the best double mutants LEIN, LDIN with the V and T substitutions seen at position 501 of the single-point mutants. Sequence LEIV has the best affinity overall (−5.1 kcal/mol).

The best quadruple mutants are included in the last column of Figure 4(b). Several mutants contain an L to Q substitution at position 455 that either maintains affinity at the same level or makes it slightly smaller; for example, sequences QEIV and QEIT have similar affinities with LEIV and LEIT, QDIV has a somewhat reduced affinity compared to LDIV.

## DISCUSSION AND CONCLUSIONS

The present study compared complexes of the human ACE2 receptor and the receptor binding domain of SARS-CoV-2 and SARS-CoV S1 proteins by all-atom MD simulations and searched for sequence substitutions at positions 455, 493, 494 and 501 of the SARS-CoV-2 S1 RBD, which might increase the virus affinity for hACE2.

Using all-atom simulations and interaction energy decomposition, we quantified contributions of the various CoV, CoV-2 and hACE2 residues on the affinities of the two complexes. Since hACE2 is negatively charged, positively charged residues are expected to favor and negatively charged residues to disfavor association. Indeed, substitutions K417/V404, K444/T431, K458/H445, G476/D463, S494/D480 inserted a positive charge or eliminated a negative charge at the CoV-2 RBD and favor the CoV-2 complex; substitutions N439/R426, L452/K439, N460/K447, E471/V458, T478/K465, E484/P471 eliminated a positive charge or inserted a negative charge and had the opposite effect.

In general, it might be expected that the accumulation of positive charge in the CoV-2 RBM would be favored, since it would increase the affinity for the negatively charged hACE2 receptor. Nevertheless, the above substitutions decrease the CoV-2 charge by one unit, relative to CoV, and have a small total effect on the relative affinity (the total electrostatic intermolecular interaction energy of these residues is −0.4 kcal/mol in favor the CoV-2 complex). The lack of accumulation of a large positive charge might help the RBMs escape recognition from other molecules than the ACE2 receptor. Furthermore, some of the above substitutions create or eliminate salt-bridge forming residues and might be favored for stability reasons. Instead, the improved recognition of hACE2 by the S1 RBM seems to rely mostly on substitutions at the interface of the complex. In the CoV-2 complex, such substitutions increasing the CoV-2 affinity are K417/V404 and F486/L473. The former introduces the salt bridge K417-D30 and the latter improves nonpolar interactions with a hydrophobic pocket consisting of hACE2 residues L79, M82 and Y83.

Residue component differences are only indicative of changes in affinity due to these substitutions. Nevertheless, many of the above conclusions are in line with a recent experimental study, which examined the impact of CoV substitutions at the CoV-2 RBM (16). Substitutions N439R, L452K, E484P and Q498Y were shown to increase and F486L to decrease the affinity of the CoV-2 complex; charge-modifying mutations E471V and S494D had an insignificant effect; thus, charge changes at some RBD positions might have a smaller impact on affinity.

Using protein computational design methods, we next examined systematically a large number of substitutions at positions 455, 493, 494 and 501 of the CoV-2 RBM, which have been evaluated as important for ACE2 recognition and cross-species transmission (13, 16). Main improvements in affinity are associated with the introduction of negative residues (E, D) at position 493 and nonpolar (I, V) or aromatic (W) at position 494; the polar CoV-2 serine can also be substituted by asparagine at that position. Insertions of a valine or the CoV threonine at position 501 yield small improvements on the affinity. The CoV-2 leucine is maintained at position 455, in sequences of high affinity; insertion of a glutamic acid at this position is only seen in conjunction with mutations at other places, without improvement in affinity.

We analyzed some complexes containing the above substitutions by 10-ns all-atom MD simulations. Figure 5(a) shows a representative simulation structure of the complex with the highest affinity triple mutant LEIV. E493 forms a stable hydrogen bond with K31, without perturbing the hotspot-31 salt bridge K31-E35. The I494 sidechain stays in proximity of the D38 sidechain (the average distance I494(CB)-D38(CB) ~ 7 Å) and forms two stable intramolecular nonpolar contacts with Y449 and L452. The I494-L452 contact replaces the structurally equivalent contacts S494-L452 and D480-K439, observed respectively in the CoV-2 and CoV RBM. The hotspot salt-bridge D38-K353 and the intermolecular salt-bridge K417-D30 are also maintained. V501 makes a non polar contact and Q498 a hydrogen bond with the K353 sidechain. L455 contacts the nonpolar moiety of K31.

**Figure 5:**
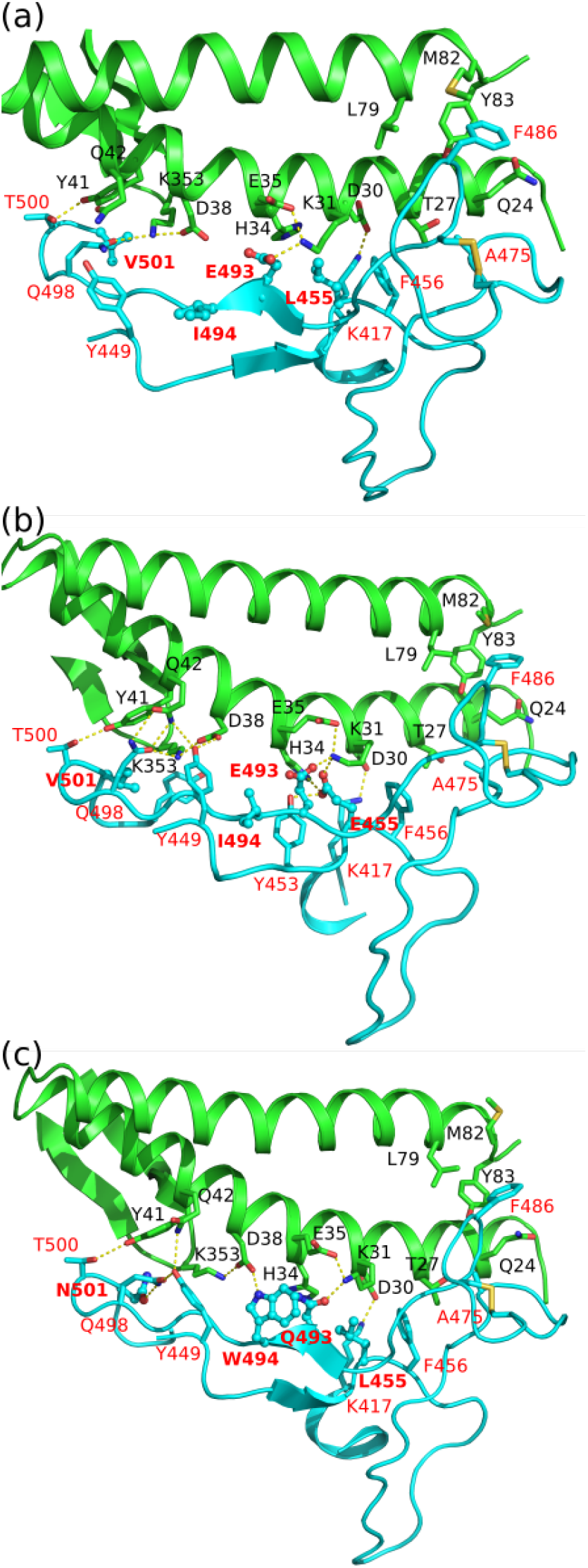
Typical MD structures of human ACE2 complex with SARS-CoV-2 designed sequences L455/E493/I494/V501 (a) E455/E493/I494/V501 (b) L455/Q493/W494/N501 (c) Substituted side chains are shown in stick-and-ball representation and red bold labels. Hydrogen bonds are indicated by dotted yellow lines.

A second interesting case is the EEIV mutant, which contains two negatively charged residues at positions 455 and 493. Figure 5(b) shows a representative simulation structure of the corresponding complex. E455 forms two stable intermolecular hydrogen bonds with K31 and H34 and an intramolecular hydrogen bond with Y453. E493 forms a second stable hydrogen bond with K31, without disrupting the salt-bridges K31-E35 and K417-D30. The I494 sidechain forms similar interactions as in the LEIN complex. The Y449 sidechain forms hydrogen bonds with D38 and Q42. The Q42 sidechain is oriented toward Q498, forming two simultaneous hydrogen bonds with it. Another hydrogen bond is formed between Q498 and K353. The hotspot-353 salt-bridge K353-D38 is also maintained.

Figure 5(c) shows a representative simulation structure of the hACE2 complex with the LQWN mutant, which contains a tryptophane substitution at position 494. The W494 sidechain is oriented towards ACE2, forming a hydrogen bond with D38.

Figure 4(c) shows the composition at positions 455, 493, 494 and 501 (CoV-2 numbering) in various viral strains encountered in other species than human (16). The Pangolin (GD) sequence composition is identical to SARS-CoV-2. Among the other sequences, only the Pangolin (GX) sequence 455L/493E/494R/501T was predicted by our design to have better affinity than CoV-2 (2.9 kcal/mol). Note though that the Pangolin (GX) RBM contains several additional substitutions with respect to CoV-2, including mutations K417V, F486L and Q498H at positions contacting hACE2. The total effect of all these substitutions might be to reduce affinity for hACE2 below the CoV-2 affinity.

At position 455, some of the coronavirus sequences of Figure 4(c), substitute L with S, R, W and Y. Among these types, an arginine is predicted in some of our designed sequences. The best sequence is REIV, with a relative affinity of −2.8 kcal/mol. Reconstruction of the designed conformation shows that R455 can form an intermolecular salt-bridge with E35 and an intramolecular salt-bridge with E493.

At position 493, the coronavirus sequences of Figure 4(c), substitute L with E, Y, K, A, R, S. Among these types, we frequently encounter a tyrosine residue in our design. The best such sequence is LYIV, with an affinity of −2.8 kcal/mol. The bat (RatG13) sequence LYRD also contains a tyrosine at this position. Our design predicts the closely related sequence LYRT, with an affinity of −0.3 kcal/mol.

At position 501, several coronavirus sequences of Figure 4(c) contain mainly small polar side chains (N,T,S) and three of the bat sequences contain a V or D residue. In our designed sequences, V, T and N are the most frequently encountered choices in sequences of high-affinity. Our design inserts a threonine residue in the best and in 12 out of the 20 top-binding sequences. In accord with this, the substitution N501T was recently shown in ref (50) to improve the CoV-2 affinity for hACE2.

In conclusion, the computational protein design calculations predict that the combinations (E, D)493 /(I, V, N, W)494 /(V, T, N)501 might increase the SARS-CoV-2 affinity for hACE2. All-atom MD simulations show that the above substitutions create new intermolecular salt-bridges or hydrogen bonds, without disrupting the hotspots 31 and 353, or the salt-bridge K417-D30. The introduction of an E sidechain at position 455 creates a hydrogen bond with H34. E or Q sidechains at position 493 form a hydrogen bonds with K31. The introduction of I, V and W sidechains at position 494 creates a nonpolar patch and promotes association; the W sidechain is positioned between D38 and E35, forming a hydrogen bond with D38. A V sidechain at position 501 makes a non polar contact with K353. Some of these interactions could be used in pharmacophore models and assist in the identification of peptidomimetic compounds and organic molecules with the potential to inhibit the binding of the viral S1 protein to hACE2.

## AUTHOR CONTRIBUTIONS

GA and SP designed the research, carried out simulations and analysis and wrote the manuscript.

## ACKNOWLEDGMENTS

SP thanks the University of Cyprus and the European Regional Development Fund and the Republic of Cyprus through the Research and Innovation Foundation (Project: INFRASTRUCTURES/1216/0060) for financial support and the Cyprus Ministry of Education, Culture, Sports and Youth for a leave of absence. The authors state no conflict of interest related to the work in this manuscript.

## REFERENCES

1. Li, F., 2014. Receptor Recognition Mechanisms of Coronaviruses: a Decade of Structural Studies. J. Virol. 89:1954–1964. https://doi.org/10.1128/jvi.02615-14.

2. Andersen, K. G., A. Rambaut, W. I. Lipkin, E. C. Holmes, and R. F. Garry, 2020. The proximal origin of SARS-CoV-2. Nat. Med. 26:450–452. https://doi.org/10.1038/s41591-020-0820-9.

3. Corman, V. M., D. Muth, D. Niemeyer, and C. Drosten, 2018. Hosts and Sources of Endemic Human Coronaviruses. In Advances in Virus Research, Elsevier, 163–188. https://doi.org/10.1016/bs.aivir.2018.01.001.

4. Wu, F., S. Zhao, B. Yu, Y.-M. Chen, W. Wang, Z.-G. Song, Y. Hu, Z.-W. Tao, J.-H. Tian, Y.-Y. Pei, M.-L. Yuan, Y.-L. Zhang, F.-H. Dai, Y. Liu, Q.-M. Wang, J.-J. Zheng, L. Xu, E. C. Holmes, and Y.-Z. Zhang, 2020. A new coronavirus associated with human respiratory disease in China. Nature 579:265–269. https://doi.org/10.1038/s41586-020-2008-3.

5. Zhou, P., X.-L. Yang, X.-G. Wang, B. Hu, L. Zhang, W. Zhang, H.-R. Si, Y. Zhu, B. Li, C.-L. Huang, H.-D. Chen, J. Chen, Y. Luo, H. Guo, R.-D. Jiang, M.-Q. Liu, Y. Chen, X.-R. Shen, X. Wang, X.-S. Zheng, K. Zhao, Q.-J. Chen, F. Deng, L.-L. Liu, B. Yan, F.-X. Zhan, Y.-Y. Wang, G.-F. Xiao, and Z.-L. Shi, 2020. A pneumonia outbreak associated with a new coronavirus of probable bat origin. Nature 579:270–273. https://doi.org/10.1038/s41586-020-2012-7.

6. Zhou, H., X. Chen, T. Hu, J. Li, H. Song, Y. Liu, P. Wang, D. Liu, J. Yang, E. C. Holmes, A. C. Hughes, Y. Bi, and W. Shi, 2020. A novel bat coronavirus reveals natural insertions at the S1/S2 cleavage site of the Spike protein and a possible recombinant origin of HCoV-19 https://doi.org/10.1101/2020.03.02.974139.

7. Li, F., 2016. Structure, Function, and Evolution of Coronavirus Spike Proteins. Annu. Rev. Virol. 3:237–261. https://doi.org/10.1146/annurev-virology-110615-042301.

8. Li, W., M. J. Moore, N. Vasilieva, J. Sui, S. K. Wong, M. A. Berne, M. Somasundaran, J. L. Sullivan, K. Luzuriaga, T. C. Greenough, H. Choe, and M. Farzan, 2003. Angiotensin-converting enzyme 2 is a functional receptor for the SARS coronavirus. Nature 426:450–454. https://doi.org/10.1038/nature02145.

9. Hoffmann, M., H. Kleine-Weber, S. Schroeder, N. Krüger, T. Herrler, S. Erichsen, T. S. Schiergens, G. Herrler, N.-H. Wu, A. Nitsche, M. A. Müller, C. Drosten, and S. Pöhlmann, 2020. SARS-CoV-2 Cell Entry Depends on ACE2 and TMPRSS2 and Is Blocked by a Clinically Proven Protease Inhibitor. Cell 181:271–280.e8. https://doi.org/10.1016/j.cell.2020.02.052.

10. Li, F., 2005. Structure of SARS Coronavirus Spike Receptor-Binding Domain Complexed with Receptor. Science 309:1864–1868. https://doi.org/10.1126/science.1116480.

11. Li, F., 2008. Structural Analysis of Major Species Barriers between Humans and Palm Civets for Severe Acute Respiratory Syndrome Coronavirus Infections. J. Virol. 82:6984–6991. https://doi.org/10.1128/jvi.00442-08.

12. Wu, K., G. Peng, M. Wilken, R. J. Geraghty, and F. Li, 2012. Mechanisms of Host Receptor Adaptation by Severe Acute Respiratory Syndrome Coronavirus. J. Biol. Chem. 287:8904–8911. https://doi.org/10.1074/jbc.m111.325803.

13. Wan, Y., J. Shang, R. Graham, R. S. Baric, and F. Li, 2020. Receptor Recognition by the Novel Coronavirus from Wuhan: an Analysis Based on Decade-Long Structural Studies of SARS Coronavirus. J. Virol. 94. https://doi.org/10.1128/jvi.00127-20.

14. Wrapp, D., N. Wang, K. S. Corbett, J. A. Goldsmith, C.-L. Hsieh, O. Abiona, B. S. Graham, and J. S. McLellan, 2020. Cryo-EM structure of the 2019-nCoV spike in the prefusion conformation. Science 367:1260–1263. https://doi.org/10.1126/science.abb2507.

15. Lan, J., J. Ge, J. Yu, S. Shan, H. Zhou, S. Fan, Q. Zhang, X. Shi, Q. Wang, L. Zhang, and X. Wang, 2020. Structure of the SARS-CoV-2 spike receptor-binding domain bound to the ACE2 receptor. Nature 581:215–220. https://doi.org/10.1038/s41586-020-2180-5.

16. Shang, J., G. Ye, K. Shi, Y. Wan, C. Luo, H. Aihara, Q. Geng, A. Auerbach, and F. Li, 2020. Structural basis of receptor recognition by SARS-CoV-2. Nature 581:221–224. https://doi.org/10.1038/s41586-020-2179-y.

17. Villa, F., N. Panel, X. Chen, and T. Simonson, 2018. Adaptive landscape flattening in amino acid sequence space for the computational design of protein:peptide binding. J. Chem. Phys. 149:072302. https://doi.org/10.1063/1.5022249.

18. Opuu, V., G. Nigro, T. Gaillard, E. Schmitt, Y. Mechulam, and T. Simonson, 2020. Adaptive landscape flattening allows the design of both enzyme: Substrate binding and catalytic power. PLOS Computational Biology 16:e1007600. https://doi.org/10.1371/journal.pcbi.1007600.

19. Mignon, D., K. Druart, S. Opuu, V. Polydorides, F. Villa, T. Gaillard, E. Michael, G. Archontis, and T. Simonson, 2020. Proteus software for physics-based protein design. J. Chem. Theory Comp..

20. Wang, F., and D. P. Landau, 2001. Efficient, Multiple-Range Random Walk Algorithm to Calculate the Density of States. Phys. Rev. Lett. 86:2050–2053. https://doi.org/10.1103/physrevlett.86.2050.

21. Olsson, M. H. M., C. R. Søndergaard, M. Rostkowski, and J. H. Jensen, 2011. PROPKA3: Consistent Treatment of Internal and Surface Residues in Empirical pKa Predictions. J. Chem. Theory Comp. 7:525–537. https://doi.org/10.1021/ct100578z.

22. Brünger, A. T., and M. Karplus, 1988. Polar hydrogen positions in proteins: Empirical energy placement and neutron diffraction comparison. Proteins 4:148–156. https://doi.org/10.1002/prot.340040208.

23. Jo, S., T. Kim, V. G. Iyer, and W. Im, 2008. CHARMM-GUI: A web-based graphical user interface for CHARMM. J. Comp. Chem. 29:1859–1865. https://doi.org/10.1002/jcc.20945.

24. Phillips, J. C., R. Braun, W. Wang, J. Gumbart, E. Tajkhorshid, E. Villa, C. Chipot, R. D. Skeel, L. Kalé, and K. Schulten, 2005. Scalable molecular dynamics with NAMD. J. Comp. Chem. 26:1781–1802. https://doi.org/10.1002/jcc.20289.

25. Martyna, G. J., D. J. Tobias, and M. L. Klein, 1994. Constant pressure molecular dynamics algorithms. J. Chem. Phys. 101:4177–4189. https://doi.org/10.1063/1.467468.

26. Feller, S. E., Y. Zhang, R. W. Pastor, and B. R. Brooks, 1995. Constant pressure molecular dynamics simulation: The Langevin piston method. J. Chem. Phys. 103:4613–4621. https://doi.org/10.1063/1.470648.

27. Huang, J., and A. D. MacKerell, 2013. CHARMM36 all-atom additive protein force field: Validation based on comparison to NMR data. J. Comp. Chem. 34:2135–2145. https://doi.org/10.1002/jcc.23354.

28. Jorgensen, W. L., J. Chandrasekhar, J. D. Madura, R. W. Impey, and M. L. Klein, 1983. Comparison of simple potential functions for simulating liquid water. J. Chem. Phys. 79:926–935. https://doi.org/10.1063/1.445869.

29. Durell, S. R., B. R. Brooks, and A. Ben-Naim, 1994. Solvent-Induced Forces between Two Hydrophilic Groups. J. Phys. Chem. 98:2198–2202. https://doi.org/10.1021/j100059a038.

30. Neria, E., S. Fischer, and M. Karplus, 1996. Simulation of activation free energies in molecular systems. J. Chem. Phys. 105:1902–1921. https://doi.org/10.1063/1.472061.

31. Essmann, U., L. Perera, M. L. Berkowitz, T. Darden, H. Lee, and L. G. Pedersen, 1995. A smooth particle mesh Ewald method. J. Chem. Phys. 103:8577–8593. https://doi.org/10.1063/1.470117.

32. Tuckerman, M., B. J. Berne, and G. J. Martyna, 1992. Reversible multiple time scale molecular dynamics. J. Chem. Phys. 97:1990–2001. https://doi.org/10.1063/1.463137.

33. Michael, E., S. Polydorides, T. Simonson, and G. Archontis, 2017. Simple models for nonpolar solvation: Parameterization and testing. J. Comp. Chem. 38:2509–2519. https://doi.org/10.1002/jcc.24910.

34. Weeks, J. D., D. Chandler, and H. C. Andersen, 1971. Role of Repulsive Forces in Determining the Equilibrium Structure of Simple Liquids. J. Chem. Phys. 54:5237–5247. https://doi.org/10.1063/1.1674820.

35. Levy, R. M., L. Y. Zhang, E. Gallicchio, and A. K. Felts, 2003. On the Nonpolar Hydration Free Energy of Proteins: Surface Area and Continuum Solvent Models for the Solute-Solvent Interaction Energy. J. Am. Chem. Soc. 125:9523–9530. https://doi.org/10.1021/ja029833a.

36. Aguilar, B., R. Shadrach, and A. V. Onufriev, 2010. Reducing the Secondary Structure Bias in the Generalized Born Model via R6 Effective Radii. J. Chem. Theory Comp. 6:3613–3630. https://doi.org/10.1021/ct100392h.

37. Gohlke, H., and D. A. Case, 2003. Converging free energy estimates: MM-PB(GB)SA studies on the protein-protein complex Ras-Raf. J. Comp. Chem. 25:238–250. https://doi.org/10.1002/jcc.10379.

38. Swanson, J. M. J., R. H. Henchman, and J. A. McCammon, 2004. Revisiting Free Energy Calculations: A Theoretical Connection to MM/PBSA and Direct Calculation of the Association Free Energy. Biophys. J. 86:67–74. https://doi.org/10.1016/s0006-3495(04)74084-9.

39. Simonson, T., T. Gaillard, D. Mignon, M. S. am Busch, A. Lopes, N. Amara, S. Polydorides, A. Sedano, K. Druart, and G. Archontis, 2013. Computational protein design: The proteus software and selected applications. J. Comp. Chem. 34:2472–2484. https://doi.org/10.1002/jcc.23418.

40. Tuffery, P., C. Etchebest, S. Hazout, and R. Lavery, 1991. A New Approach to the Rapid Determination of Protein Side Chain Conformations. J. Biom. Struct. Dyn. 8:1267–1289. https://doi.org/10.1080/07391102.1991.10507882.

41. Still, W. C., A. Tempczyk, R. C. Hawley, and T. Hendrickson, 1990. Semianalytical treatment of solvation for molecular mechanics and dynamics. J. Am. Chem. Soc. 112:6127–6129. https://doi.org/10.1021/ja00172a038.

42. Hawkins, G. D., C. J. Cramer, and D. G. Truhlar, 1995. Pairwise solute descreening of solute charges from a dielectric medium. Chem. Phys. Lett. 246:122–129. https://doi.org/10.1016/0009-2614(95)01082-k.

43. Schaefer, M., and M. Karplus, 1996. A Comprehensive Analytical Treatment of Continuum Electrostatics. J. Phys. Chem. 100:1578–1599. https://doi.org/10.1021/jp9521621.

44. Polydorides, S., N. Amara, C. Aubard, P. Plateau, T. Simonson, and G. Archontis, 2011. Computational protein design with a generalized born solvent model: Application to asparaginyl-tRNA synthetase. Proteins 79:3448–3468. https://doi.org/10.1002/prot.23042.

45. Polydorides, S., and T. Simonson, 2013. Monte carlo simulations of proteins at constant pH with generalized born solvent, flexible sidechains, and an effective dielectric boundary. J. Comp. Chem. 34:2742–2756. https://doi.org/10.1002/jcc.23450.

46. Gaillard, T., and T. Simonson, 2014. Pairwise decomposition of an MMGBSA energy function for computational protein design. J. Comp. Chem. 35:1371–1387. https://doi.org/10.1002/jcc.23637.

47. Metropolis, N., A. W. Rosenbluth, M. N. Rosenbluth, A. H. Teller, and E. Teller, 1953. Equation of State Calculations by Fast Computing Machines. J. Chem. Phys. 21:1087–1092. https://doi.org/10.1063/1.1699114.

48. Yan, R., Y. Zhang, Y. Li, L. Xia, Y. Guo, and Q. Zhou, 2020. Structural basis for the recognition of SARS-CoV-2 by full-length human ACE2. Science 367:1444–1448. https://doi.org/10.1126/science.abb2762.

49. Li, X., E. E. Giorgi, M. H. Marichannegowda, B. Foley, C. Xiao, X.-P. Kong, Y. Chen, S. Gnanakaran, B. Korber, and F. Gao, 2020. Emergence of SARS-CoV-2 through recombination and strong purifying selection. Sci. Adv. 6:eabb9153. https://doi.org/10.1126/sciadv.abb9153.

50. Yi, C., X. Sun, J. Ye, L. Ding, M. Liu, Z. Yang, X. Lu, Y. Zhang, L. Ma, W. Gu, A. Qu, J. Xu, Z. Shi, Z. Ling, and B. Sun, 2020. Key residues of the receptor binding motif in the spike protein of SARS-CoV-2 that interact with ACE2 and neutralizing antibodies. Cell. Mol. Immunol. 17:621–630. https://doi.org/10.1038/s41423-020-0458-z.

